# Illuminating the complete ß-cell mass of the human pancreas - signifying a new view on the islets of Langerhans

**DOI:** 10.1101/2023.11.15.566244

**Authors:** Joakim Lehrstrand, Wayne I. L. Davies, Max Hahn, Olle Korsgren, Tomas Alanentalo, Ulf Ahlgren

## Abstract

Pancreatic islets of Langerhans play a pivotal role in regulating blood glucose homeostasis, but critical information regarding their mass, distribution and composition is lacking within a whole organ context. Here, a new 3D imaging pipeline was applied to generate a first complete account of the insulin-producing islets throughout the human pancreas at a microscopic resolution and within a maintained spatial 3D context. These data show that human islets are far more heterogenous than previously accounted for with regards to their size distribution and cellular make up. By deep tissue 3D imaging, this in-depth study demonstrates that 50% of the human insulin-expressing islets are virtually devoid of glucagon-producing a-cells, an observation with significant implications for both experimental and clinical research.

**One Sentence Summary:** New islet heterogeneity identified in the human pancreas

## MAIN

By virtue of their roles in regulating blood glucose homeostasis, the hormone producing islets of Langerhans have been the subject of intense research for well over a century ^1^. Still, understanding their distribution and intra-islet organization across the human pancreas remains limited. In humans, estimates of islet numbers, volumes and cellular ratios vary significantly. Inter-individual biodiversity apart, this may be attributed to differences in selected analytical methods, most of which rely on extrapolation of a limited amount of two-dimensional (2D) data, or on subfractions of isolated islets. Hence, the non-diabetic (ND) human pancreas has been reported to contain between 1-14.8 million islets, with a mean diameter of ∼108 µm and a total islet volume of 0.5-2.0 cm^3^. Similarly, analyses of the relative contributions of different endocrine cell-types to overall islet composition display significant differences (^2–5^ and references therein). A consensus of several reports suggest that human islets are composed of ∼60% insulin (INS)-producing ß-cells and 30% glucagon (GCG)-producing α-cells. The remaining 10% primarily consists of somatostatin-producing 8-cells, followed by pancreatic polypeptide (PP) and ghrelin-producing PP-cells and ε-cells, respectively (^2^ and references therein). With regards to overall distribution, the islet mass is suggested to be both unevenly ^6^ and evenly ^7^ distributed within the pancreas.

Islet cells require intercellular communication to function properly, including a large number of paracrine signals that act within islets together with exogenous neural inputs to ensure correct islet responses to glucose and other metabolites ^8,9^. It is generally acknowledged that diabetes is a disease that involves all islet cell-types, not only ß-cells ^10^. Therefore, a precise understanding of islet organization throughout the entire organ (including size, distribution and hormonal composition) is critical to fully appreciate the significance of islet architecture and whole organ pancreatic distribution for normal physiology and disease etiology. In this study, mesoscopic optical 3D imaging approaches were used to provide the first whole organ account of the β-cell mass distribution (i.e., islet volume and number, as well as 3D spatial location) across the entire volume of the human pancreas. Data presented here provides convincing evidence for a previously unknown heterogeneity in islet composition and demonstrates that as much as 50% of human islets contain only a few (<1%) or no glucagon-producing α-cells. Apart from the direct implications for pre-clinical and clinical research, these observations will serve as a foundation for the generation of precise anatomical atlases of the human pancreatic endocrine system and how it is affected under pathological conditions.

## RESULTS

### The first 3D reconstruction of the complete ß-cell distribution of the human pancreas

Recently, the labeling and imaging of cm^3^-volumes of human pancreatic tissue at a microscopic resolution was demonstrated, while maintaining the 3D context of all labeled cells ^11^. Implementing adapted protocols (see methods), the entire β-cell distribution throughout the pancreas of a deceased ND donor was analyzed (see **Extended Data Table 1**). Briefly, donated pancreata were divided into “discs”, using a 3D printed matrix, (**Fig. 1A** and **Extended Data Fig. 1**) that were labeled for insulin and scanned individually by near infrared optical projection tomography (NIR-OPT) ^12,13^ (**Fig. 1B** and **C**, **Extended Data Fig. 2, Movie 1**). Selected regions of interest (ROIs) were also scanned by higher resolution light sheet fluorescence microscopy (LSFM). For NIR-OPT data, an INS^+^ object in this study is defined as a distinct body of cells stained for insulin that cannot be separated from each other at the resolution observed during segmentation (here, OPT scan settings results in an isotropic resolution of ∼21 µm). By aligning the resultant tomographic datasets together in 3D space, an entire pancreas could be “rebuilt” with regards to the 3D distribution of INS^+^ cells (**Fig. 1D, Extended Data Movie 2** and **Fig. 3**). Segmentation of INS^+^ signals allowed for the extraction of a complete picture of all INS^+^ objects including their individual volumes and spatial 3D coordinates throughout the volume of the pancreas. Hereby, a range of comprehensive statistical assessments of the β-cell mass (BCM), or more precisely β-cell volume, distribution could be extracted within a maintained spatial 3D context that is not dependent on the extrapolation of partially sampled data.

**Fig. 1.**
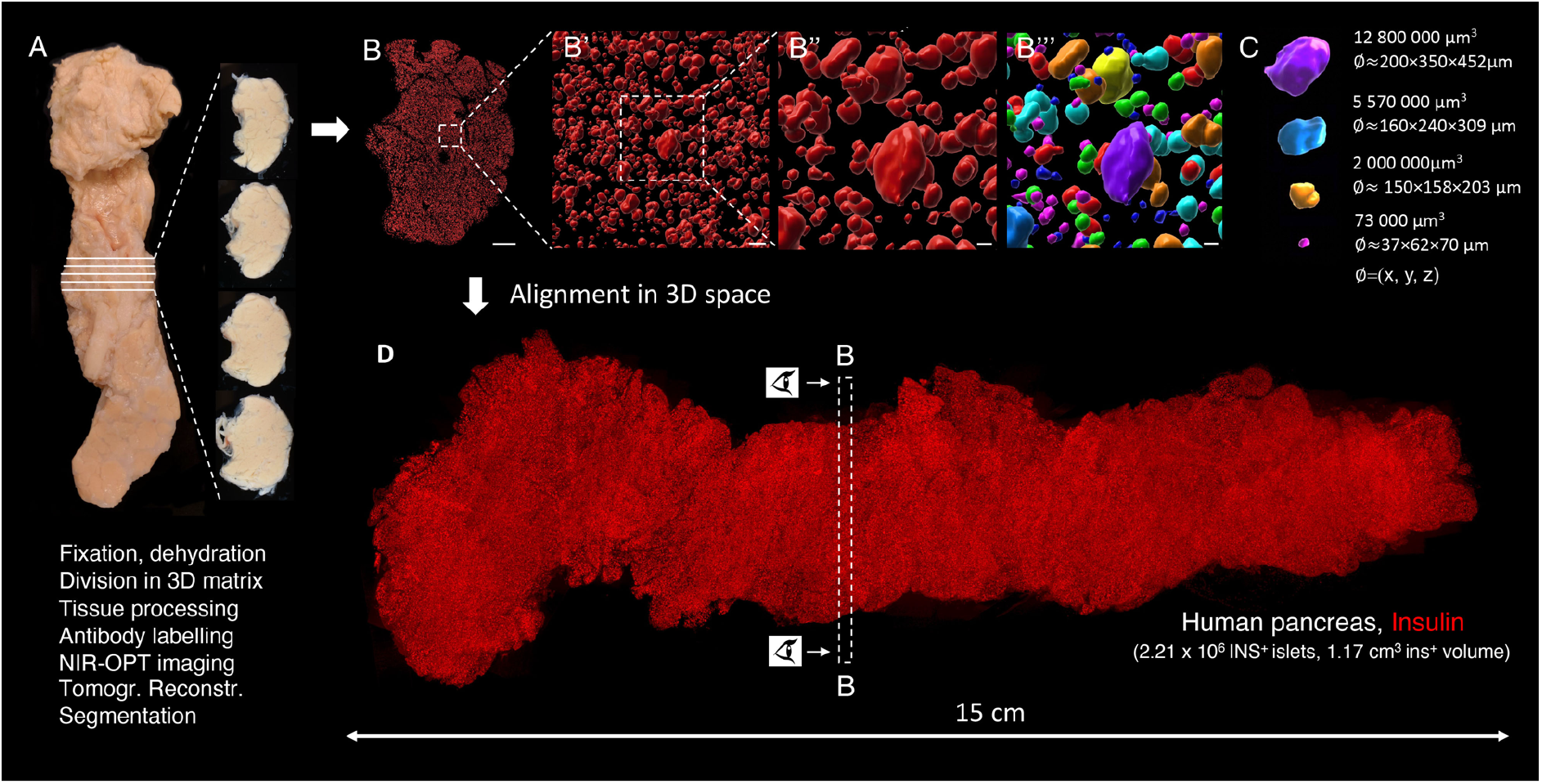
Generation of volumetric and 3D-spatial data sets of the complete β-cell distribution across the human pancreas. The intact pancreas (**A**) was fixed directly after procurement from a deceased donor (H2457, see **Extended Data Table 1**) and dehydrated in 100% ethanol. Using a 3D printed matrix, the pancreas was divided into 2.5 mm discs, with known spatial origins, allowing for penetration of tissue processing chemicals and antibodies. Each disc was scanned individually by NIR-OPT (see **Extended Data Movie 1**) or LSFM before offline segmentation and 3D rendering of individual islet β-cell volumes (**B, C**). In **B’’’**, each pseudo-colored object corresponds to a specific β-cell volume with known 3D coordinates. (**D**) By combining individual tomographic datasets in 3D space, new data sets could be created that encompass every insulin-stained object: in this case there were 2.21 x 10^6^ INS^+^ objects with a total volume of 1.17 cm^3^, here displayed as a single 3D maximum intensity projection (see also **Extended Data Movie 2** and **Fig. 3**). Eye symbols in **D** illustrates the angle of view of disc shown in **B**. Scale bar in B is 3000μm in B, 200 μm in B’ and 70 μm in B’’ and B’’’.

### Revealing the normal distribution of the human β-cell mass - an absolute volumetric assessment

The ND donor pancreas displayed in **Fig. 1** (H2457, see **Extended Data Table 1**) comprises an INS^+^ volume of 1.17 x 10^12^ µm^3^ (=1.17 cm^3^) that consists of 2.21 x 10^6^ separate INS^+^ islets at the applied resolution. To regionally refine these values, the pancreas was divided into four regions consisting of the head (region 1, using the indentation of the superior mesenteric vein as a boundary) and regions 2-4 consisting of portions 1/3 in length of the remainder of the pancreas. These analyzes showed that the ß-cell density is relatively uniform across the length (head to tail) of the human pancreas (**Fig. 2A** and **Fig. 2B**), although variations exist between individual discs, possibly due to regional differences in vascular and ductal densities (**Fig. 2A**).

**Fig. 2.**
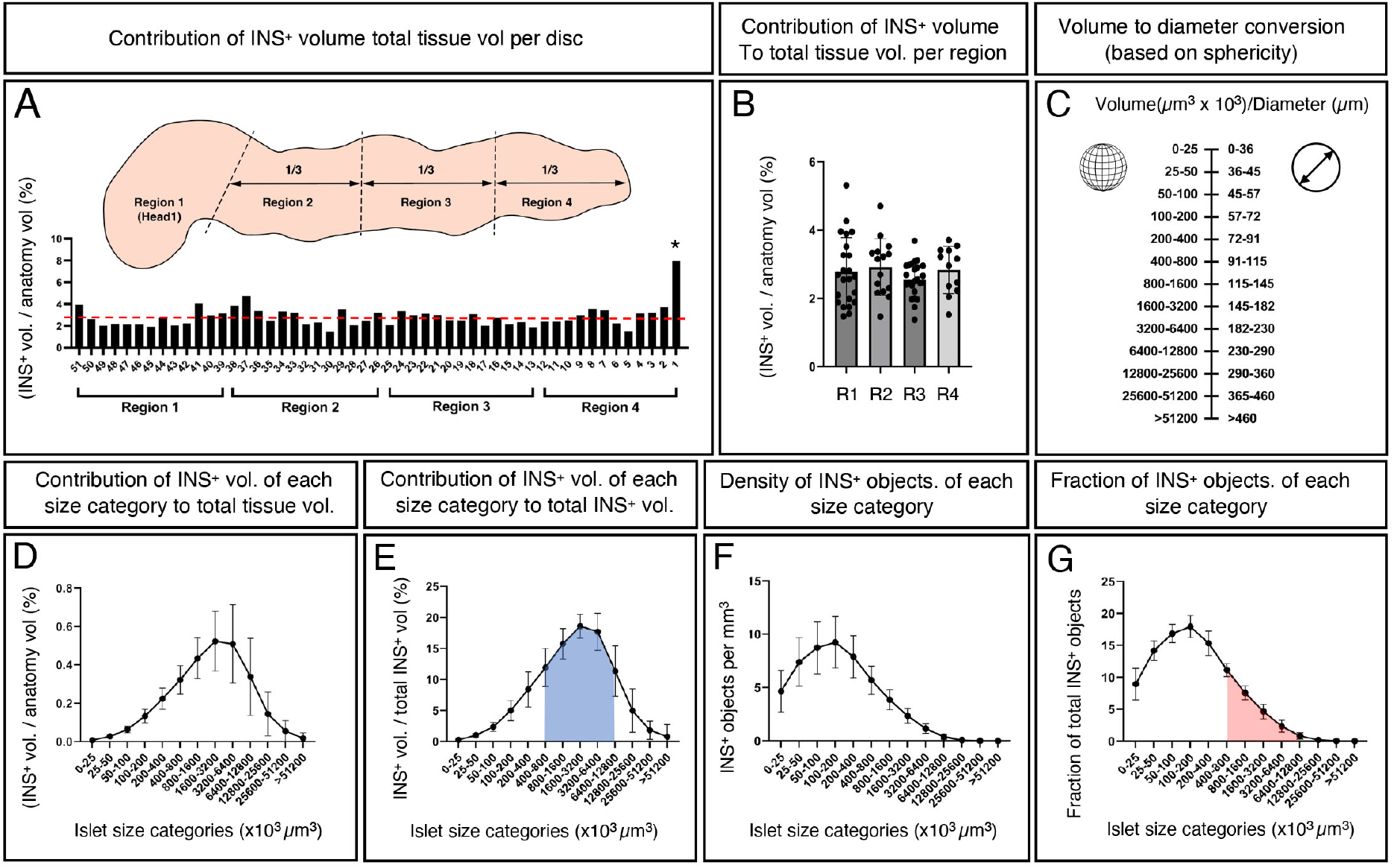
3D quantification and statistical assessment of the complete human pancreatic β-cell distribution. (**A**) INS^+^ volume normalized to the tissue volume for each disc (H2457, see **Extended Data Table 1**). Note, whereas the walls of vessels and ducts are included into the total tissue volume, “empty space” contained within these structures are not. The average INS^+^ density across the entire pancreas is 2.8% (red broken line). Schematic inset illustrates the regional separation of the pancreas into four defined regions. (**B**) A pooled bar graph showing the same data as in **A** redefined for each of the four regions of the human pancreas. (**C**) A graph illustrating the corresponding diameter of spherical objects for each size category (for comparison to 2D stereological studies), noting that islet sphericity is not perfect and this value is not the same as the average 3D diameter (see **Extended Data Fig. 3**). (**D-G**) Graphs showing the volume of each INS^+^ islet size category represented as (**D**) volumetric density normalized to the entire tissue volume, (**E**) volumetric density normalized to the entire INS^+^ volume, (**F**) number of INS^+^ objects per mm^3^ per size category and (**G**) the fraction of INS^+^ objects per INS^+^ islet size category (**G**). The blue area in **E** corresponds to the red area in **G**, showing that the majority of the BCV (∼75%) consists of only ∼25% of islets. *Indicates a small tissue disc at the very tip of the tail that encompassed only 573 INS^+^ objects (see **Extended Data Fig. 1**). Graphpad robust regression and outlier removal (ROUT) with Q=1% identified this small sample as an outlier and it is was removed from subsequent analysis. Error bars show mean ± SD.

As islet size has important functional implications (see discussion below), the 2.21 x 10^6^ measured INS^+^ objects were subdivided into different size categories, where the fraction that each size category represented as a volume normalized to the entire tissue volume was calculated (**Fig. 2D**). To better understand islet size distribution, further calculations were performed that included the volume fraction out of total INS^+^ volume (**Fig. 2E**), INS^+^ objects per size category per mm^3^ (**Fig. 2F**) and the fraction of INS^+^ objects constituted by each size category (**Fig. 2G**). Similar data were obtained from all regions (1-4), alluding to a relatively uniform size distribution of INS^+^ islets across the length of the human pancreas (**Extended Data Fig. 4**). To simplify comparisons of these “absolute” 3D volumetric measurements to previous morphometric 2D studies, each volume size category was converted to the diameter of a spherical object for a particular corresponding range interval (**Fig. 2C**). Note, however, that islets are not perfectly spherical objects (See **Extended Data Fig. 5** for sphericity analyzes of iso-surfaced volumes). Presented data show that around 75% of the total BCV consists of INS^+^ islets in the range of 400 x 10^3^ – 12800 x 10^3^ μm^3^ (which corresponds to spheres ∼91 - 290 μm in diameter), but that this volume only corresponds to ∼26 % of all INS^+^ islets (see blue and red color-coding in **Fig. 2E** and **Fig. 2G**, respectively). Further, when all pancreatic islets are considered, these data indicate that the average islet diameter (based on insulin staining in 3D space) decreases to ∼65 μm (or ∼75 μm when accounting for typical tissue processing shrinkage and assuming perfect sphericity (i.e., Ψ=1)). Of note, it was previously demonstrated that at the current resolution, using OPT, the average 3D-diameter of islets, based on the insulin signal, deviated less than ± 5% compared to the diameter measured on Hematoxylin/Eosin stained tissue sections of the same islets ^11^.

### Human and mouse islets have a similar contribution of islet size categories to the overall BCM but differ in their organization within the organ

As demonstrated optical by 3D imaging, rodent islets are heterogeneously distributed both within and between the three primary, splenic, duodenal and gastric, pancreatic lobes ^14–16^, and analyses of different diabetes disease models showed that islets of different sizes can be unequally affected during disease progression ^17–20^. E.g., in the Non Obese Diabetic (NOD) model for naturally induced T1D, smaller islets are more susceptible to autoimmune destruction, whereas large centrally located islets even appear to possess a compensatory growth potential ^17^. However, similar types of assessments of the human pancreas have been effectively hindered by technological limitations, and our knowledge about the normal distribution of the human β-cell mass in 3D space is limited.

By defining small medium and large sized islets as making up 1/3 each of the total β-cell mass of the entire pancreas (**H2457, Extended Data Fig. 3** and **Table 1**) and cross reference this information back to the 3D reconstruction of each disc we could not detect any pronounced heterogeneities in the distribution of islets of different size-categories within the organ (**Fig 3A, B, Extended Data Fig 6** and **Movie 3**). Possibly, larger islets could appear less frequent in the absolute periphery of the organ, but this observation was not consistent across all samples/regions (**Fig. S6**). Whereas the relative contribution of each size category to the overall BCM is very similar between mice and humans (compare **Fig. 3B** and **D**), the spatial organization of islets of different size categories differ, and in analogy with previous reports (see above) large mouse islets were preferentially located in vicinity of the organ’s central axis, following the main pancreatic duct (Compare **Fig. 3B, D, Extended Data Movie 3** and **4**). Heat maps of the human samples, visualizing the average distance of each islet to its 5 nearest neighbors within each size category, confirmed this picture (**Fig. 3E-G**) and in contrast to previous stereological assessments we could find convincing evidence for islet routes ^7^ or varying islets densities in specific areas of the pancreas, head to tail ^6^. The average distance to the 5 nearest neighbors was longest for the large islets and shortest for the smallest, reflecting the relative number of islets within these size categories (**Fig. 3H, I**).

**Fig. 3.**
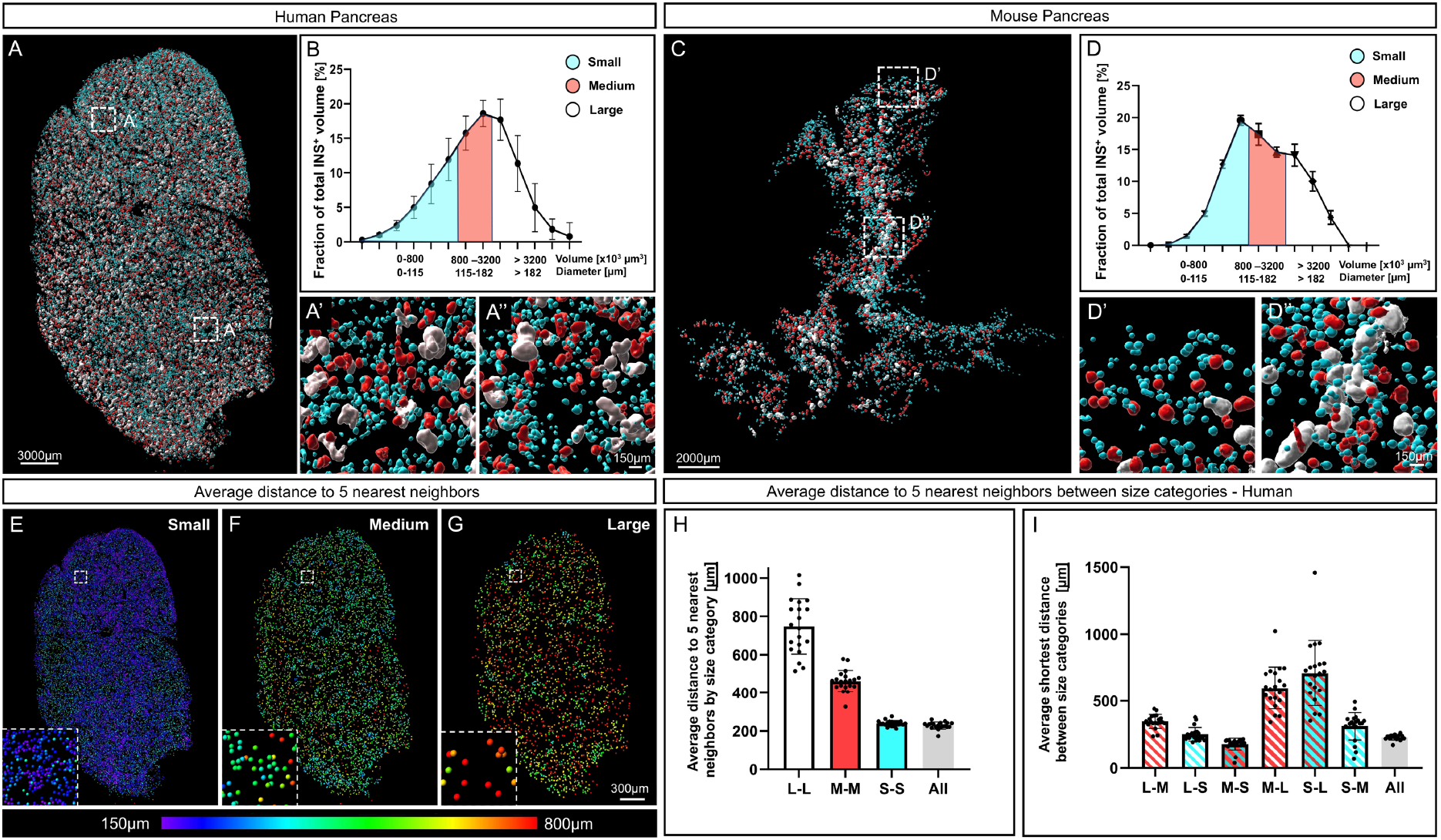
Spatial arrangement and relative contribution of islet size categories in man and mice. (**A**) Image of a representative pancreatic disc in which the insulin signal has been segmented and pseudo colored according to size. Each size category (blue, small; red, medium; white, large) corresponds to 1/3 of the total β-cell volume (**B**) of the entire pancreas depicted in **Extended Data** Fig 3, incorporating 2.21 x 10^6^ INS^+^ islets. (**C**) Image of a representative mouse pancreas pseudo colored as in the same way (**A**) based on all INS^+^ islets in C57BL/6 pancreata at 10 weeks (n=5, including 28108 INS^+^ islets). (**E-G**) Heat maps showing the average distance to the 5 nearest objects for each size category in the disc depicted in (**A**). (**H, I**) Bar graphs showing the average distance to the 5 nearest neighboring objects (islets) by size category (**H**) and between size categories (**I**) calculated from one disc from each region (1-4) in five donor pancreata (H2456, H2457, H2466, H2506 and H2522, see **Extended Data Table 1**). Whereas the relative contribution of each size category to the overall BCM is very similar between the two species, the larger islets are predominantly located close to the center axis of the organ in mice. In contrast human islets of all size categories are more evenly distributed in the organ (see also **Extended Data Fig. 6**).

Altogether, this first global assessment of human pancreatic β-cells at the current level of resolution provides strong evidence that the BCM is relatively evenly distributed along the pancreas from head to tail, that the average islet size is significantly smaller than what has been reported by 2D stereological studies or by assessments of isolated islets, and the human β-cell mass organization differ significantly from that of the mouse. Further, they open for precise 3D dimensional, combined volumetric, assessments of specifically antibody targeted cells and structures throughout the volume of human donor pancreata in different pathological settings.

### The majority of the human INS^+^ islets are devoid of GCG expressing cells

It is generally accepted that a direct interaction between α-cells and β-cells is crucial for glycemia management ^21^. As a first step in exploring this relationship in the entire human pancreas, 3D OPT-based imaging approaches for large scale studies of intra-islet cellular composition were applied to representative tissue discs from regions 1-4 from ND pancreata encompassing both genders and an age range of 20-45 years of age (n=5, see **Extended Data Table 1**). 3D maximum intensity projection (MIP) views (**Extended Data Figs. 7 and 8**), covering the entire depth of tens of thousands of islets for each scan, revealed a dramatic degree of heterogeneity in intra-islet composition that is not consistent with the consensus 2:1 ß-cell to α-cell ratio. Strikingly, a very high proportion of human islets, primarily in the size range below 800 x 10^3^ μm^3^ in volume (≈90 μm in diameter), appeared to be completely devoid of GCG altogether or possess just a few α-cells (**Extended Data Fig. 7 D-E**). To confirm this observation, and to resolve the possibility that individual GCG^+^ cells were not included at the applied resolution when using OPT (∼21 µm), LSFM scans of ROIs within the same tissue discs at a spatial resolution of 1.8 μm were generated. Hereby, cellular ratios across the volume of the investigated tissue discs at a high cellular resolution could be determined (**Fig. 4 A-E**, **Extended Data Movie 5** and **Fig. 9**). Analyzes of two ROIs from each of the four anatomical regions in five different ND pancreata (see **Extended Data Table 1**) revealed that on an average 50.2 % of the INS^+^ islets contain <1% GCG-expressing (GCG^+^) cells. These islets, however, although abundant in numbers, corresponded to only ∼16% of the total islet volume (red bars in **Fig. 4F-G**). Serial 2D sections stained for INS and GCG, including every section of individually assessed islets identified by LSFM imaging, supported these important new observations of the presence of a significant number of INS^+^GCG^−^ islets in the human pancreas **(Extended Data Fig. 10)**. Specifically, these data indicate that most GCG^+^ cells reside within larger islets (>800 x 10^3^ μm^3^ in volume; >115 μm in diameter) and that the absolute majority of the GCG^+^ volume is comprised by islets containing 1-30% GCG^+^ cells, with 10-20% GCG^+^ cells making the greatest contribution (**Fig. 4 I-J).** Altogether, these studies provide evidence for a dramatic heterogeneity in islet size and composition in the human pancreas, with a majority of islets consisting of INS^+^, but not GCG^+^ cells.

**Fig. 4.**
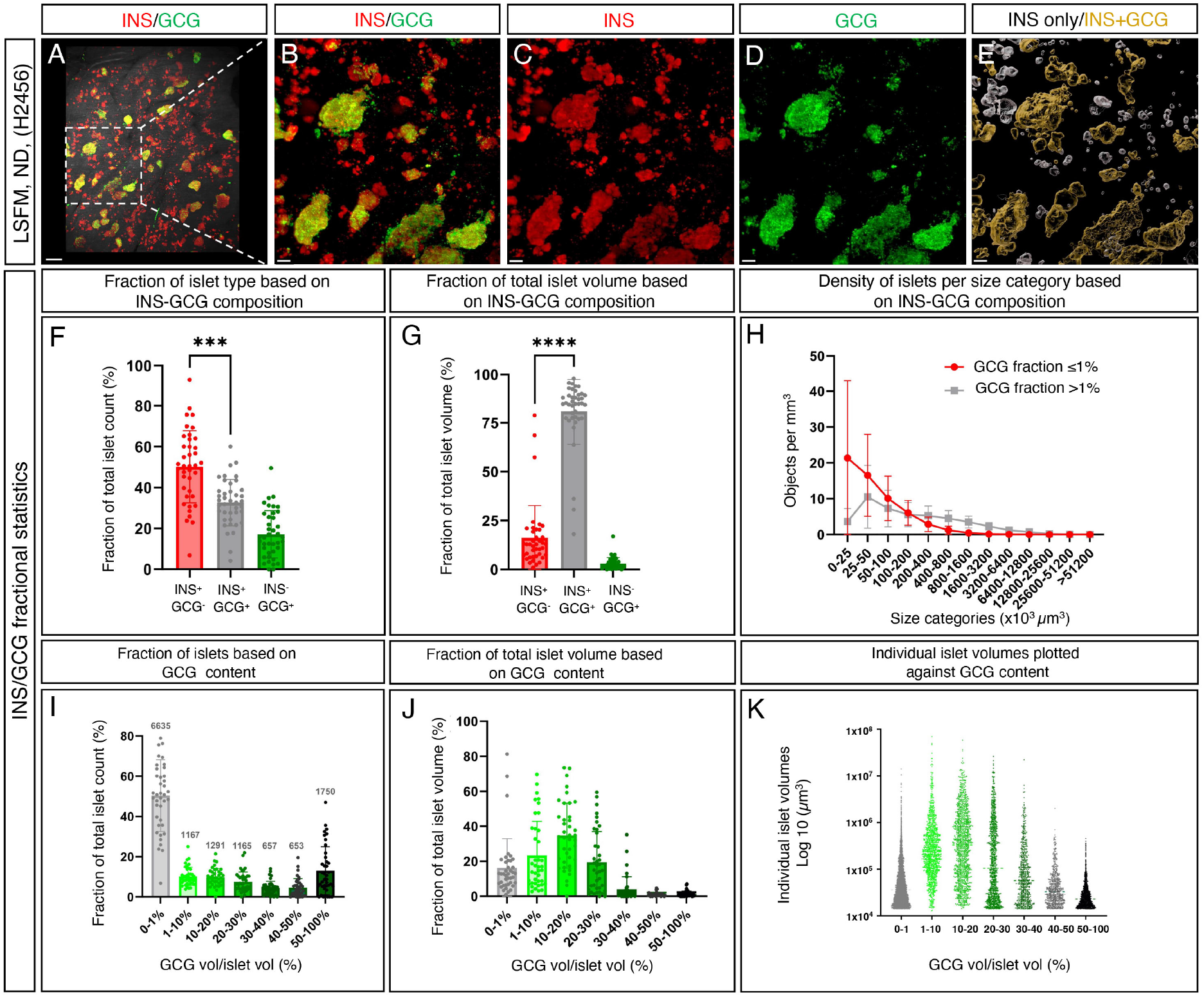
High resolution deep tissue 3D imaging reveals striking heterogeneities in human islet endocrine cell composition. (**A-D**) 3D Maximum intensity projection (MIP) from a LSFM scan of a representative ROI (see methods) from a ND donor pancreas stained for insulin (INS, red) and glucagon (GCG, green) showing the presence of INS^+^GCG^−^ islets. (**E**) Pseudo-colored transparent surface reconstructions where islets with <1% GCG^+^ volume are white and islets with >1% GCG^+^ cells are orange (see also **Extended Data Movie 5)**. (**F**) A graph showing the contribution of islets with < 1% GCG^+^ volume to the total number of INS^+^GCG^−^, INS^−^GCG^+^ and INS^+^GCG^+^ islets. (**G**) A graph showing the contribution of islets with < 1% GCG^+^ volume to the total volume of INS- and GCG-expressing cells. (**H**) Densities of INS^+^GCG^−^ (red line) and INS^−^GCG^+^ and INS^+^GCG^+^ (grey line) in **F** and **G** represented as a function of size category. (**I-K**) GCG volume/ islet volume as fraction of total islet count per ROI (I), as fraction out of total islet volume (J) and the individual islet volumes (K) respectively. Data are derived from ROIs from regions 1-4 from five different ND donor pancreata (see **Extended Data Table 1 and Fig. 7**) scanned at 1.8 μm spatial resolution. Scale bar in **A** is 200 μm and scale bars in **B**-**E** are 100 μm. *** and **** donates a statistical significance of p ≤ 0.001 and p ≤ 0.0001 respectively.

## DISCUSSION

Already in 1906, Lydia Dewitt stated “no organ or tissue of the body has been the subject of more thought or investigation than have the areas of Langerhans, especially during the past years” ^22^. Given the ongoing diabetes pandemic and the immense efforts still active in both basic and clinical diabetes arenas, this statement still holds true over a century later. This present report provides the first in depth whole organ account of the complete islet ß-cell mass distribution within the human pancreas, while maintaining the spatial 3D context (i.e., volume, shape and location) of every single islet. Assuming that every islet contains β-cells expressing insulin (INS^+^), this study has detected and analyzed at least 2.21 x 10^6^ INS^+^ islets in a representative healthy ND donor pancreas. Unlike earlier studies, these measurements are not dependent on extrapolation of a limited amount of 2D or 3D data and suggest that BCV is uniformly distributed across the pancreas, from head to tail. Further, this study demonstrates that the average human islet is significantly smaller than previously reported (see ^23^ and references therein), which has important sampling and functional implications as discussed below. It has previously been suggested that small islets contain a relatively large number of INS^+^ cells (63%) compared to large islets where the number of INS+ cells is lower (39%), and that these smaller islets have greater insulin secretion ^24^. Strikingly, the deep tissue imaging approach applied in this report, which covers the entire volume of hundreds of thousands of islets, show that very few islets display the literature consensus composition of around 60% INS^+^ and 30% GCG^+^ cells. Instead, the human endocrine pancreas comprises ∼50% INS^+^ islets that belong to the smaller end of the spectrum of islet size ranges (still larger than 29 μm in diameter), which are essentially devoid of GCG^+^ cells. Noteworthy, the investigated pancreata contained between ∼50.2 and 63% INS^+^GCG^−^ islets, with the exception of H2506 which contained “only” ∼30% INS^+^GCG^−^ islets (**Extended Data Fig. 11 and Table 1)**, still a considerable figure. The dramatic number of INS^+^GCG^−^ islets observed here raises important questions about islet specialization and functionality. A large number of paracrine signals have been demonstrated to act within islets to ensure their proper functional responses to glucose and other metabolites ^25^. Further, studies imply that heterologous contact between α-cells and ß-cells are crucial for glycemia management ^21^. Most significantly, insulin (INS) together with other signaling molecules inhibit glucagon (GCG) secretion, whereas glucagon stimulates insulin secretion. Cholinergic innervation to human islets is sparse ^26^. However, α-cells have been reported to provide ß-cells with acetylcholine, which enhances insulin release by binding to muscarinic receptors ^26^, and it has been shown that blocking GCG signaling in islets limits insulin secretion ^27^. As such, it is very surprising that the vast number of INS^+^ islets devoid of GCG (i.e., INS^+^GCG^−^ islets) reported here has not been recognized previously. This may be due, until now, to the lack of an accurate and sensitive 3D imaging perspective, which allows for the simultaneous study of multiple islets throughout a large tissue depth. It is also possible that a general assumption that islets have a typical mixed cell population has eluded their discovery and subsequent reporting in most 2D stereological studies. For example, when assessing cryosection or wax embedded samples with islets lacking GCG^+^-cells, it has been plausible to assume that GCG^+^-cells would be present on a consecutive slide or that the observed islet was not “representative”. Whereas isolated islets have become an invaluable tool to understand islet biology and diabetes ^28^, data presented here emphasize that human islet isolates do not reflect the full range of islets sizes and cellular compositions in the endogenous islet population. In addition, islets distributed for research are commonly associated with cultivation artefacts ^29^. This is of particular significance since several *in vitro* studies suggest that smaller islets have superior cellular function compared with larger islets (See ^30^ and references therein). Interestingly, it has also been demonstrated that small mouse islets after isolation are virtually devoid of α-cells ^31^, but perhaps this subset of murine islets actually represents the subpopulation of human INS^+^GCG^−^ islets identified in this present study.

Whereas insulin is key for glucose metabolism, glucagon primarily controls amino acid metabolism ^32^. The observed heterogeneity in islet composition suggests that pancreatic islets of different cellular arrangements might promote specialized functional niches in order to meet different metabolic demands. For example, it is possible that there is an endogenous plasticity in islet constitution in response to prolonged exposure to specific metabolites. As such, it is plausible that ß-cells within islets with different proportions of hormonal cells may have different metabolic activities. Further studies incorporating other hormonal cell-types and markers for different cellular subtypes may shed light on these questions, as well as to the possibility that islets of different configurations and putative functional roles are distinctly innervated.

Altogether, the results presented here advocate for a re-evaluation of how an islet of Langerhans is defined. Further, they strongly suggest that islet size and cellular composition should be carefully considered in any research or clinical setting aiming at reflecting or reproducing the endogenous islet population of the human pancreas. As such, these data are likely to greatly impact a broad range of undertakings in islet and pancreas research, including *ex vivo* experimentation of isolated, strategies for islet generation of islets by stem-cell or bio-printing approaches, the optimization of clinical islet transplantation protocols that better reflect the endogenous microcellular environment of the pancreas and many more. The developed possibility to study the pancreas for the first time from all angles and through its entire depth (with known 3D coordinates, volumes and shapes for all labeled objects) will provide a unique opportunity to identify features of normal pancreatic anatomy and disease pathologies that would otherwise be extremely challenging to recognize using other techniques. Finally, the developed approach applied in this study was able to overcome the frequent issue of reagent penetration in optical imaging of larger tissues. As such, the methods presented here should be translatable to a plethora of studies that target a variety of other human and non-human tissues, potentially in any physiological and/or pathophysiological context, with essentially a full freedom of target selectivity.

## MATERIALS AND METHODS

### Ethics declaration

All work involving human tissue was conducted in accordance with the Declaration of Helsinki ^33^ and in the European Council’s Convention on Human Rights and Biomedicine ^34^. Consent for organ donation for use in research was obtained from the donor prior to death via the Swedish National Donor Registry (https://www.socialstyrelsen.se/en/apply-and-register/join-the-swedish-national-donor-register/) or from relatives of the deceased donors conferred by the donor’s physician and documented in their medical records. The study was approved by the Regional Ethics Committee in Uppsala, Sweden (Dnr 2017/1471-32, 2023-01845-01).

### Organ isolation and processing

Pancreata were obtained after declared brain death from 5 multi-organ donors registered within the framework for the Nordic Network for Clinical Islet Transplantation (NNCIT), with clinical data relevant to this study listed in Extended Data Table 1. The donated pancreata were dissected from the duodenum in Ringer’s acetate solution (Braun Melsungen, Germany) and washed in 1 × PBS (Medicago, Sweden) before fixation in 4% formaldehyde (Solveco, Sweden). After 24 h, the fixative was replaced with fresh 4% formaldehyde solution and fixed for another 24 h. Fixed samples were then stepwise dehydrated in ethanol: twice with 75% (v/v) ethanol in H_2_O and twice with 96% (v/v) (VWR, Sweden) at 4°C. At this stage, pancreata were stored and transported in 96% (v/v) ethanol at room temperature (RT). In order to remove residual fat, pancreata were extensively washed with 96% (v/v) ethanol at 4 °C on an orbital shaker, replacing with fresh ethanol solution daily until the alcohol became transparent (i.e. no excess fat droplets present in the samples). Subsequently, all organs were imaged using a Nikon D5200 camera (see **Extended Data Fig. 1**).

### Tissue preparation for 3D imaging

A slicing matrix was designed with a section thickness of 2.5 mm, providing an optimal balance between reagent penetration and resolution/magnification to distinguish individual insulin-positive (INS^+^) objects for thresholding and quantification (Tinkercad, Autodesk, USA), using a 3D printer (Prusa i3 MK3S, Prusa Research, Czech Republic). Pancreata were mounted in 1.5% (w/v) low melting temperature agarose (Cat. No. 50100; Lonza, USA) at 37°C in the custom-made 3D-printed matrix (Fig. S1). Subsequently, whole pancreata were cut into 2.5 mm thick “slabs” and the agarose carefully removed. Tissue slabs wider than 2 cm were cut in two to ensure a good fit in the field of view when applying near infra-red optical projection tomography (NIR-OPT) microscopy (see below). Images were taken during the slicing process to document X, Y and Z coordinates for each slab (**Extended Data Fig. 1**). Tissue slabs were then stored in 100% methanol (MeOH; Cat. No. 67-89-4; Fisher Scientific, Sweden) at −20 °C until further processing.

### Whole-mount immunohistochemistry staining and tissue clearing

Tissue slabs were initially taken from −20°C to −80°C in 100% MeOH for at least 1 h, then cycled between −80°C to RT at least 5 times (typically for 2-2.5 h per step) to increase antibody penetration. To bleach pigmented cells and reduce autofluorescence, each sample was incubated for 12 h (or overnight) in bleaching solution containing hydrogen peroxide solution (H_2_O_2_; Cat. No. H1009; Sigma-Aldrich, Merck, Germany) at a final concentration of 15% (v/v)), dimethyl sulfoxide (DMSO; Cat. No. D5879; Sigma-Aldrich, Merck, Germany) and MeOH (H_2_O_2_:DMSO:MeOH) in a ratio of 3:1:2. After a further 8-12 h incubation in fresh bleaching solution, samples were washed twice in 100% MeOH at 30-60 min per wash at RT.

For whole-mount immunolabeling, all samples were rehydrated stepwise (33%, 66% and 100%, 1 h per step) from MeOH/TBST (0.15 M NaCl, 0.1 M Tris-HCl and 0.1% Triton® X-100 (Cat. No. 108603; Sigma-Aldrich, Merck, Germany), pH 7.5), followed by a further wash with 100% TBST for 1 h. Next, slabs were incubated at 37°C in blocking solution comprising TBST supplemented with 10% heat-inactivated goat serum (Cat. No. CL1200-500; Cedarlane, Canada), 5% DMSO (Cat. No. D5879; Sigma-Aldrich, Merck, Germany) and 0.01% sodium azide (NaN_3_) for 2 days. After blocking, samples were incubated for 7 days at 37°C with primary antibodies diluted in blocking solution after first being filtered through a 25 mm Acrodisc® 0.45 μm syringe filter (Cat. No. 4614; Pall Corporation, USA) to remove any potential artifacts such as aggregated antibody complexes. Primary antibodies used consisted of guinea pig anti-insulin/pro-insulin (INS; Cat. No. 16049; Progen Biotechnik, Germany; diluted 1:3000) and rabbit anti-glucagon/pro-glucagon (GCG; Cat. No. HPA036761; Atlas Antibodies, Stockholm, Sweden; diluted 1:1000). Following incubation with primary antibodies, all samples were rigorously washed three times in TBST heated to 37°C on a rotator for 1 h per wash step, then overnight, followed by two washes the next day in TBST as before. Next, samples were incubated at 37°C for 5 days with filtered secondary antibodies in blocking solution, including donkey anti-guinea pig IRDye^®^ 680 (Cat. No. 926-68077; Li-Cor Biosciences, USA; diluted 1:250) and donkey anti-rabbit Alexa Fluor^®^ 594 (Cat. No. 711-585-152; Jackson ImmunoResearch, UK; diluted 1:500). Once again, antibody solutions were filtered to remove any potential fluorophore precipitates. After secondary labelling, all slabs were washed three times in TBST on a rotator (at 37°C; 1 h per wash step, then overnight), followed by two washes the next day in TBST.

For clearing and subsequent imaging, slabs were mounted in 1.5% low melting point agarose at ∼37°C, cooled to set at RT for ∼30 min, then moved to 4°C to completely solidify for at least 3-4 h, preferably overnight. All mounted tissue slabs were cut to remove excess agarose to form clean, straight edges without any acute angles and subsequently dehydrated in aqueous MeOH solutions, moving stepwise from 33%, to 66% and finally 100% (1 h per step). Mounted samples were then dehydrated through at least 5 cycles of 100% MeOH (to remove all traces of H_2_O) with each step conducted overnight with gently shaking. When fully dehydrated, slabs were optically cleared using a 1:2 mixture of benzyl alcohol (Cat. No. 109626; VWR, Sweden) and benzyl benzoate (BABB) (Cat. No. 105860010; Acros Organics, USA), which was replaced every 12-24 h at least 3-5 times until optimally cleared before imaging.

### Near infra-red optical projection tomography (NIR-OPT) and light sheet fluorescence microscopy (LSFM) imaging

Mounted human pancreatic slabs were scanned submerged in BABB as previously described ^11,35^ in a custom-built near-infrared optical projection tomography (NIR-OPT) scanner ^12^, constructed using a Leica MZFLIII stereomicroscope (Leica, Germany) with a CoolLED pE-4000 LED fluorescence light source (Ludesco Microscoped, USA) and an Andor iKon-M (Andor Technology, UK) camera with a tilted mirror and a step motor for sample rotation. A zoom factor of 1.25× was used for all samples, rendering an isotropic voxel size of ∼21 µm. Filter sets (with stated excitation (Ex) and emission (Em) parameters) used were as follows: insulin (INS), Ex: HQ 665/45 nm, Em: HQ 725/50 nm; glucagon (GCG), Ex: 565/30nm, Em: 620/60nm; and anatomy (autofluorescence, AF), Ex: 425/60, Em: 480 nm LP.

To verify the presence of glucagon negative (GCG^−^) islets, selected pancreatic slabs stained for INS and GCG, and previously scanned by NIR-OPT, were reimaged at higher resolution using an UltraMicroscope II (Miltenyi Biotec, Germany), which comprised a 1× Olympus objective (Olympus PLAPO 2XC) coupled to an Olympus MVX10 zoom body, a 3000 step chromatic correction motor and a lens corrected dipping cap MVPLAPO 2× DC DBE objective. Samples were scanned at 1.6× magnification with a numerical aperture of 0.141, a depth of 1000 µm with a 5 µm stepsize, giving a voxel size of 1.9 × 1.9 × 5 µm, with a dynamic focus set to 10 images across the field of view. Light sheets were merged using the built-in projection function. Filter sets used were as follows: INS, Ex: 650/45, Em: 750/60; GCG, Ex: 580/25, Em: 625/30; and anatomy (AF), Ex: 470/40, Em: 525/50, where an exposure time for all channels was kept at 300 ms. The resultant datasets were saved in *ome.tif format native to ImSpectorPro software (version 7.0.124.9; LaVision Biotex, Germany), before being converted into 3D projection *ims files using Imaris File Converter software (version 10.0.0; Bitplane, UK).

### Post-NIR-OPT image pre-processing, 3D reconstruction and surfacing

For post-NIR-OPT processing and 3D reconstruction, all generated images were processed using the same pipeline as follows: (i) in order to increase the signal to noise (S:N) ratio within each image, the pixel range of acquired NIR-OPT frontal projections was adjusted individually to display minima and maxima; (ii) a contrast limited adaptive histogram equalization (CLAHE) algorithm was applied to equalize large contrast differences in immunofluorescence ^15^, with a tile size of 48 by 48 with sharpening for all signal channels; (iii) the axis of rotation was centred computationally post-scanning using a Discrete Fourier Transform Alignment algorithm (DFTA) from the open-access DSP-OPT software package ^36,37^ (https://github.com/ARDISDataset/DSPOPT); (iv) processed datasets were reconstructed to tomographic sections using NRecon software (version 1.7.0.4, SkyScan Bruker microCT, Belgium) with added misalignment compensation and ring artifact reduction; (v) all resultant *.bmp images were converted to *.ims file format as before using Imaris File Converter software.

Using Imaris software (version 10.0.0; Bitplane, UK), 3D surfaces of reconstructed OPT scans for anatomy (AF), INS^+^ and GCG^+^ signal were generated using a built-in automated batch processing pipeline with manually adjusted threshold levels where needed, applying both a gaussian blur and background subtraction (largest object: 158 µm) both to reduce noise, including objects with a voxel value above 5. To correct for the signal gradient in insulin staining, a two-step surfacing method was applied. Firstly, a surface to effectively segment peripheral high intensity objects was generated. Secondly, an additional surface aiming to segment low intensity objects was added and included a filter to exclude overlapping objects (with a minimum 10 µm^3^ overlap) compared to the original segmentation. The two resultant 3D volumes were then merged to account for objects across an intensity gradient (**Extended Data Fig. 2**). Finally, artifacts outside the tissue volume (determined by AF) were manually excluded.

Quality control of the surfacing method was determined by manually counting islets containing β-cells in 3 regions of interest (ROIs) for 3 slabs (H2457), in ImageJ (version 1.53k, National Institutes of Health, USA) for signal versus final segmentation, taking into consideration 7069 islets in the analyzes (see **Extended Data Fig. 2** and **Extended Data Table 2**). Accuracy was evaluated as a percentage of segmentation count vs signal count. This surfacing approach showed high levels of conformity to manual counting; therefore, the pipeline was applied to the analyzes of the whole pancreas.

### 3D alignment of the whole human pancreas

To fully reconstruct the whole human pancreas (see **Fig. 1 and Extended Data Fig. 3**), individual tissue slab 3D *ims datasets were imported and manually aligned in Imaris 3D space. Specifically, each slab was sequentially orientated based on the original anatomical images, where autofluorescent large ducts and vessels were used to guide accurate alignment, until the whole pancreas consisting of 51 tissue slabs was completely reconstructed.

### Assessment of insulin islet 3D distribution and organization

To determine islet spatial organization in 3D space, the INS^+^ surfaces were first divided into three size categories (Fig 3.) in Imaris, corresponding to equal fractions of the total INS^+^ volume. Surfaces were then converted into spots by first applying the distance transformation function which effectively generates a channel based on the surface volumes. The resulting channels werethen the basis for generating spots that more accurately correspond to the initial surface segmentation. The spots function was used to evaluate average distance to neighboring objects within categories, as well as determining the average shortest distance between size categories. In the spot function, distances are calculated between the center of mass of any two objects. A similar pipeline for comparison between human and mouse INS^+^ islet distribution was carried out on five 10-week-old male C57BL/6 J control mice from a pre-processed and segmented public dataset ^18^. Division into size categories was performed as for the human dataset.

### Determining insulin and glucagon composition in OPT and LSFM

To evaluate the frequency of GCG^−^ islets in OPT scans of human pancreatic slabs (**Extended Data Fig. 7 and 8**), 3D surfaces were generated using Imaris software of the total number of INS^+^ and GCG^+^ objects in 4 slabs (one taken from each of the four pancreatic regions) from 5 donors as described above. In addition, to generate surfaces representing all islets (independent of INS and GCG composition), INS^+^ and GCG^+^ surfaces were duplicated as GCG^+^ objects were seemingly larger. Therefore, surfacing GCG^+^ more accurately reflect the islet border. In this case, all INS^+^ surfaces were subjected to an overlap exclusion filter of a minimum of 10 µm^3^ overlap with GCG^+^ volumes. The two resulting surfaces were then merged to represent all islets. Sub-surfaces were generated from this collection of total islet 3D volumes depending on the ratio overlap of INS^+^ and GCG^+^ surfaces; these were defined into three classes as follows: (i) INS^+^GCG^+^, (ii) INS^+^GCG^−^, and (iii) INS^−^GCG^+^, where negative surface signal was defined as being less than 1% of immunolabelling for any given endocrine cell-type to account for background noise.

To determine islet INS and GCG ratios in LSFM scans (**Fig. 4** and **Extended Data Fig. 9**), two types of surfaces were similarly generated using Imaris software, one aimed at distinguishing the islet border and the other to segment INS^+^ and GCG^+^ cells, respectively (**Extended Data Fig. 13**). These analyzes were performed on ROIs sized 1.7 × 1.7 × 1 mm. As part of the pre-processing pipeline, all channels were subjected to layer normalization in Imaris software. Surfaces of stained INS^+^ and GCG^+^ objects were generated with an added background subtraction of 10 µm and exclusion of all objects below 3 × 10^3^ µm^3^ to remove noise. Moreover, to effectively segment the border of pancreatic islets, INS and GCG channels were surfaced using absolute intensity with a smoothing texture detail 3.78 µm, where the two resultant surfaces were merged to better define the entire islet (**Extended Data Fig. 13**). Further, an exclusion filter was applied for objects <1.4 × 10^4^ µm^3^ to only include objects with a minimum diameter of 29 µm (assuming sphericity) from the analyzes therefore excluding single cells from the analysis.

### Post 3D imaging histology and validation of glucagon negative islets

Slabs derived from different regions of the pancreas stained for insulin and glucagon, then 3D scanned with OPT followed by LSFM, were washed several times in 100% MeOH to remove all traces of BABB, followed by stepwise rehydration in a series of 70%, 50%, 30% and 10% (v/v) ethanol to 1 × PBS for 1 h at RT with gentle shaking per step. Agarose was removed by first washing the slabs in 0.29 M sucrose (Cat. No. 10319003; Fisher Scientific, Sweden) in 1 × PBS at 57°C, then two further washes with 0.29 M sucrose solution at RT with careful manual removal of agarose where required. After agarose removal, tissues were cryoprotected by incubating in 30% (w/v) sucrose in 1 × PBS overnight at 4°C to prevent the formation of ice crystals during the freezing process, embedded and snap frozen in NEG-50 (Cat. No. 11912365; Fisher Scientific, Sweden), and stored at −80°C. 10 um thick sections were collected onto SuperFrost Plus glass slides (Cat. No. 10149870; Fisher Scientific, Sweden), air dried at RT and washed in TBST for 10 min. Tissue sections were blocked in 10% fetal bovine serum (FBS; Cat. No. 11550356; Sigma-Aldrich, Merck, Germany) for 1 h at RT and re-stained with insulin (diluted 1:10000) and glucagon (diluted 1:5000) in blocking solution at RT overnight. Slides were washed 3 x 5 min in TBST and incubated with 4′,6-diamidino-2-phenylindole (DAPI) and the following secondary antibodies in blocking solution for 2 h at RT: Alexa Fluor 488^®^ goat anti-guinea pig IgG H&L (Cat. No. ab150185; Abcam, UK; diluted 1:500 and Alexa Fluor 594^®^ donkey anti-rabbit IgG H&L (Cat. No. 711-585-152; Jackson ImmunoResearch, UK; diluted 1:500). After incubation, slides were washed in 3 x 5 min in TBST and mounted with Vectashield^®^ mounting medium (Cat. No. H-1000; Vector Laboratories, USA).

All 2D sections were scanned using the automated Axio Scan.Z1 Slide Scanner (ZEISS, Germany), equipped with a Colibri 5/7 light source. DAPI, INS^+^ and GCG^+^ cells were imaged using an Axiocam 506 microscope camera (ZEISS, Germany) with a Plan-Apochromat 20x/0.8 M27 objective and the following filters: DAPI, 90 HE DAPI (Ex 353, Em 465); INS, 90 HE GFP (Ex 493, Em 517); and GCG, 64 HE mPlum (Ex 590, Em 618). For each islet, 3 Z-slices were taken at a range of 7 µm, with scans being saved in *.czi format and analyzed using ZEN (blue edition) microscopy software (version 3.7.97; ZEISS, Germany). To display GCG^−^ islets (**Extended Data Fig. 10**), ROIs were selected in ZEN and Z-stack images presented as orthogonal projections (maximum) with deblurring (strength, 0.5; BlurRadius, 15; and sharpness, 0).

### Statistics and reproducibility

Statistical data for surfaces generated using Imaris software from OPT and LSFM scans, including volumes, sphericity and overlap ratio between surface volumes, were exported as Excel (Microsoft Office 365, version 2304) *.xml files for quantification. For a given object, a 3D diameter was calculated as an average of the diameters in X, Y and Z axes. Graphs were generated and statistical tests performed using GraphPad Prism software (version 10.0.2; GraphPad Software, USA). Where appropriate, when selecting statistical tests, a Shapiro-Wilk test for normality was applied for each column. If the normality (gaussian distribution) test was passed, column data were subjected to a paired t-test for comparison (e.g., see **Fig. 4 F** and **Fig. 4 F**) otherwise a Wilcoxon rank test was used (**Fig. 4 G**). P-values are reported as: P > 0.05 (ns), P ≤ 0.05 (*), P ≤ 0.01 (**), P ≤ 0.001 (***), P ≤ 0.0001 (****).

## Data and code availability

Information, including raw and processed imaging datasets acquired by NIR-OPT and LSFM for all samples, are available from the corresponding author upon reasonable request. Custom generated scripts used for processing OPT data including COM-AR^37^ (alignment of axis of rotation during OPT scan setup), DFTA^37^ (alignment of axis of rotation post-OPT scanning) and CLAHE^15^ (improving islet segmentation) is compiled as a software package (together with video instructions on their implementation) at GitHub, Link “https://github.com/ARDISDataset/DSPOPT”.

## ACKNOWLEDGEMENTS

The authors thank Mrs. S. Ingvast, Mr. F. Morini and Mr. A. Ahlgren for initial technical assistance and Dr. S.M.A. Willekens for helpful comments on the manuscript. This work was supported by the Kempe Foundation, the Swedish Research Council (2017-01307, 2019-01415), an Umeå University Biotech Grant (FS 2.1.6-2026-20), the Swedish Childhood Diabetes Foundation (Barndiabetesfonden), The Novo Nordisk Foundation (NNF21OC0069771, NNF20OC0063600) and the Diabetes Wellness Foundation Sweden (PG21-6566), the Ernfors Family Fund, Nils Eric Holmstens forskningsstiftelse and Diabetesfonden.

## CONTRIBUTIONS

J.L. performed OPT, LSFM and Axioscan imaging, performed statistical analyzes, was involved in study design and the development of the computational processing pipeline. W.D. was involved in study design and performed whole mount 3D immunohistochemistry. M.H. was involved in study design and initial development of the computational processing pipeline. O.K. conceived the study together with U.A., was involved in study design and responsible for sample collection. T.A. was involved in study design and performed 2D immunohistochemistry. U.A. conceived the study and supervised the entirety of the project. All authors contributed to the interpretation of data, manuscript writing and approved the final version of the manuscript.

## Notes

### Competing Interest Statement

The authors have declared no competing interest.

