## Supplemental figures 1-2, Supplemental tables 1 & 2 for "Illuminating the complete ß-cell mass of the human pancreas - signifying a new view on the islets of Langerhans"

Joakim Lehrstrand *et al.*

### The PDF file includes:

Extended Data Figs. 1 to 13
Extended Data Table 1 and 2

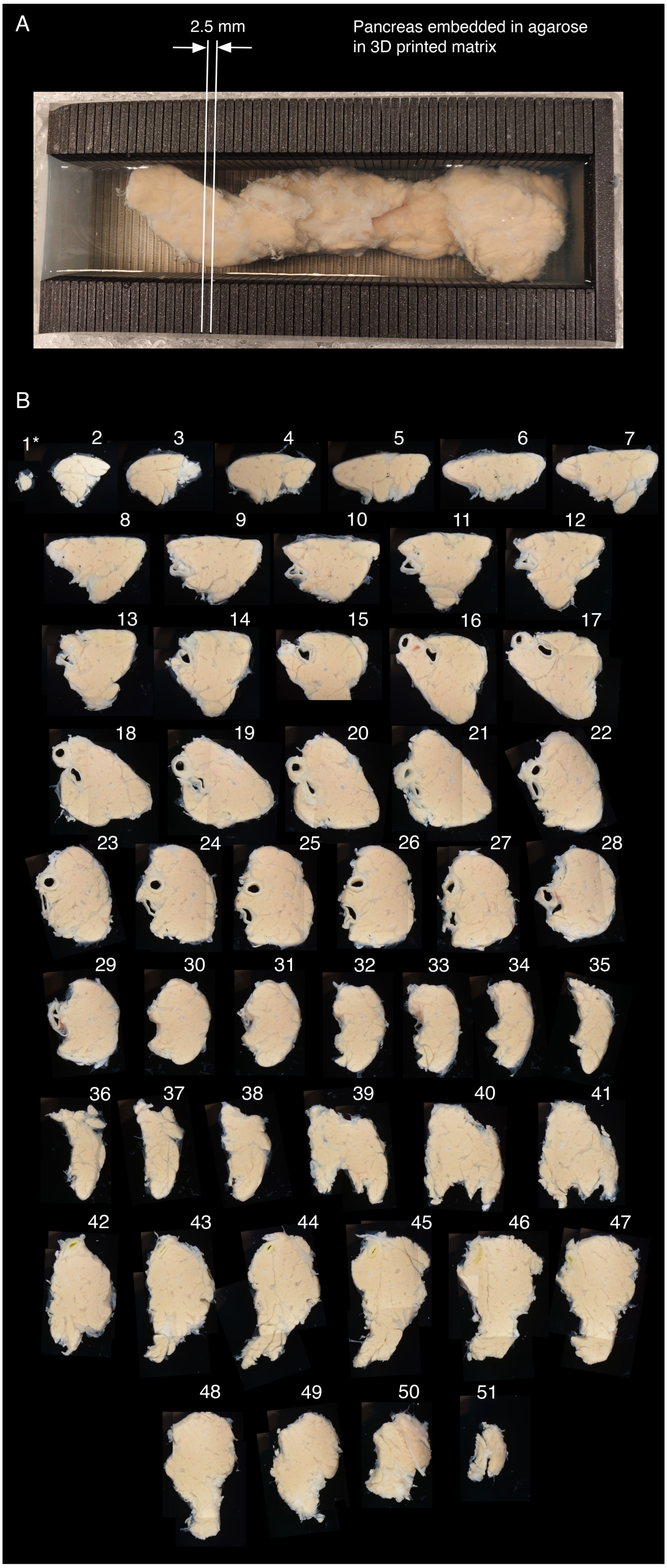


**Extended Data Fig. 1. Slicing and documentation of pancreatic discs for labeling and 3D analyzes.** (**A**) Human pancreas embedded in an agarose support in a 3D-printed matrix (grid size 2.5 mm) The opening in the matrix (left side) makes it possible to easily remove and collect each disc. (**B**) Photomicrographs of individual discs (1-51), from tail to head regions obtained from human pancreas (e.g., H2457). *Denotes a small disc containing only 547 islets (see outlier in **Fig. 2**).

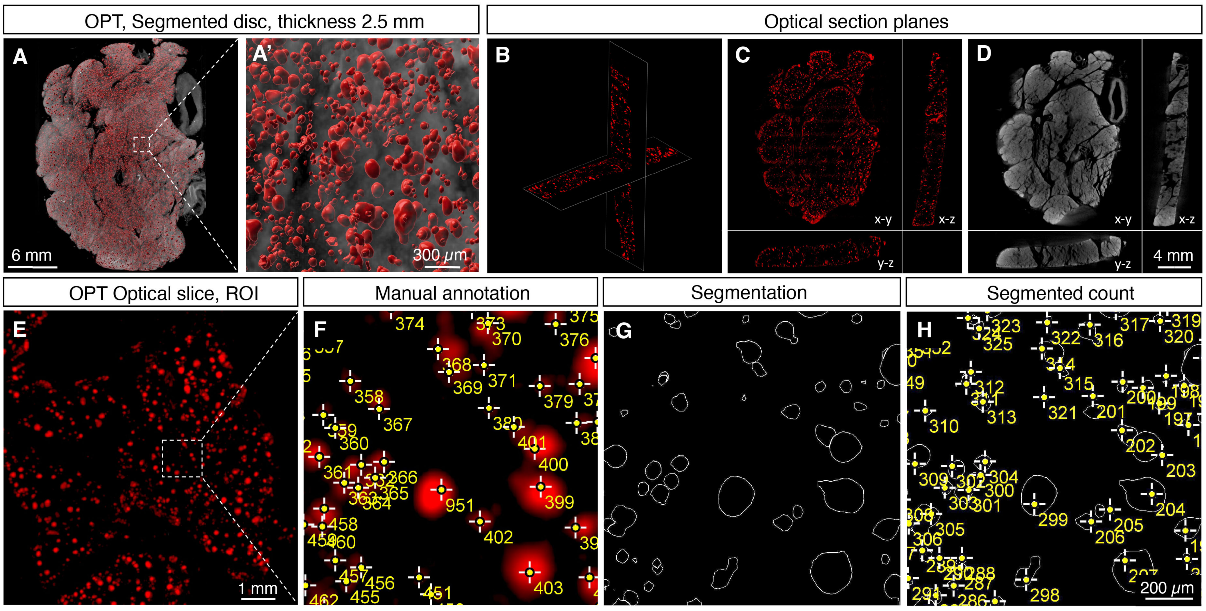


**Extended Data Fig. 2. Validation of antibody penetration and segmentation.** (**A**) Segmented INS$^+$ signal from OPT scan. (**B-D**) Optical section planes of the sample seen in panel **A** showing INS$^+$ signal throughout the volume of the tissue (**B-C**) and of the autofluorescence (AF) signal (**D**). (**E-H**) An optical slice showing exemplifying validation of segmentation, with the INS$^+$ signal observed in an optical slice ROI. Manual annotation is exemplified in panel **F**, which corresponds to the dotted box in panel **E.** The result of the segmentation pipeline is shown in panel **G** and its annotation in panel **H**. A segmentation accuracy of 7069 INS$^+$ manually annotated islets resulted in a segmentation accuracy of 102% (see **Extended Data Table 2**).

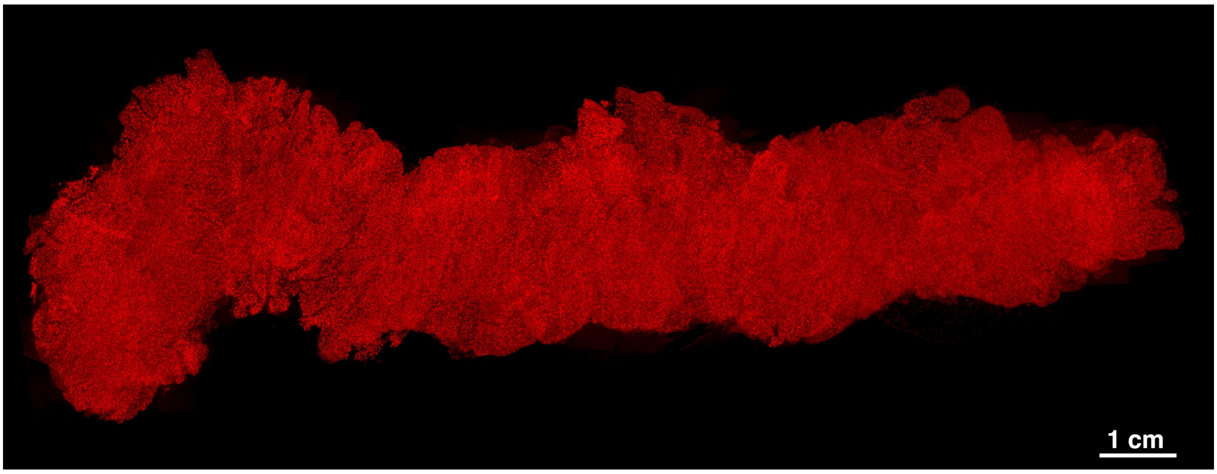


**Extended Data Fig. 3. Combined NIR-OPT datasets displaying the complete β-cell mass distribution of a representative human pancreas.** The displayed pancreas (H2457) contains 1.17 $cm^3$ $INS^+$ cells comprising 2.21 x $10^6$ $INS^+$ islets. See also **Extended data Movie 2**. Scale bar is 1 cm.

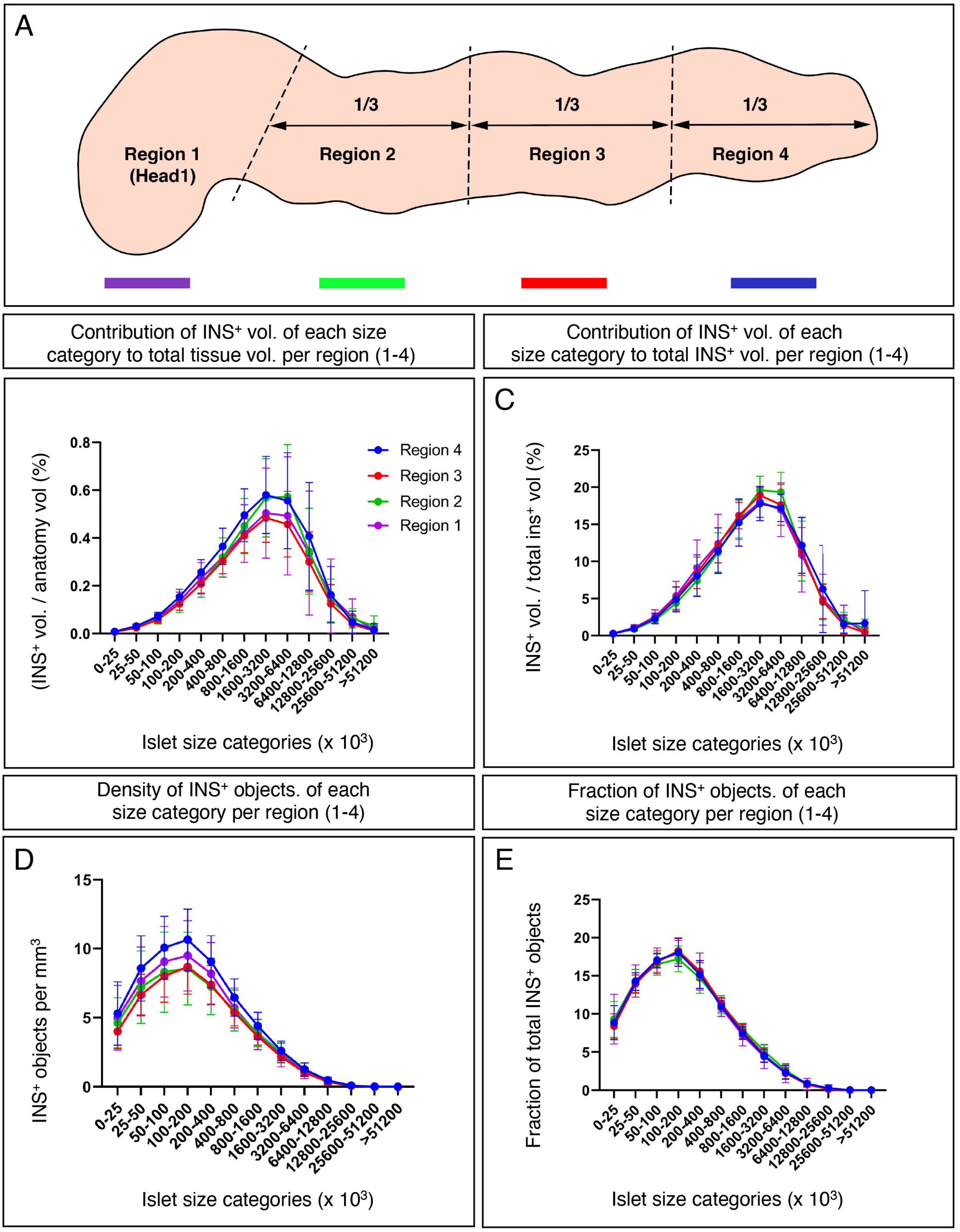


**Extended Data Fig. 4. Statistical analyzes of the size distribution of $INS^+$ objects per pancreatic region (1-4).** (**A**) Schematic illustration of regional division of the pancreas (see main text for details). (**C-E**) For each region, graphs showing the volume of each $INS^+$ islet size category represented as (i) normalized to the entire tissue volume of each disc (**B**), (ii) normalized to the entire $INS^+$ volume (**C**), (iii) the density per $mm^3$ per $INS^+$ islet size category (**D**) and (iv) the fraction of $INS^+$ objects per $INS^+$ islet size category (**E**). Error bars show mean ± SDE.

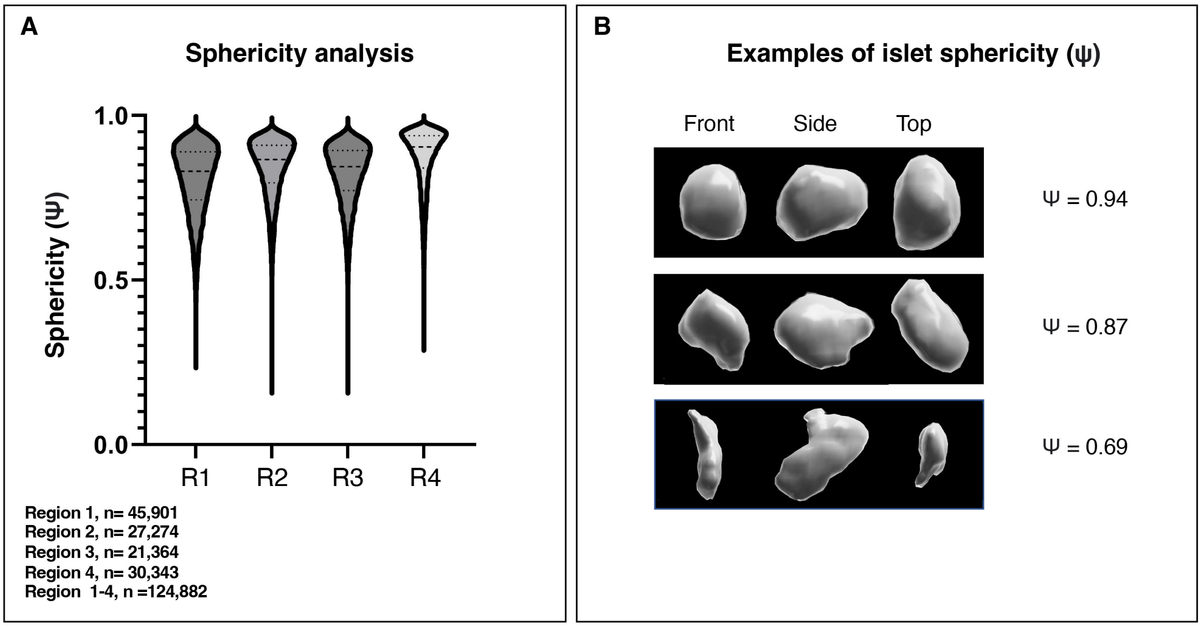


**Extended Data Fig. 5. Analysis of islet sphericity. (A)** Violin plots showing islet sphericity and the number of islets covered by the analysis for each region (1-4), where average sphericity (Ψ) was calculated to have a value of 0.83 for a total of 124,882 $INS^+$ islets derived from H2457. (**B**) Examples of segmented islet volumes and their corresponding sphericity values.

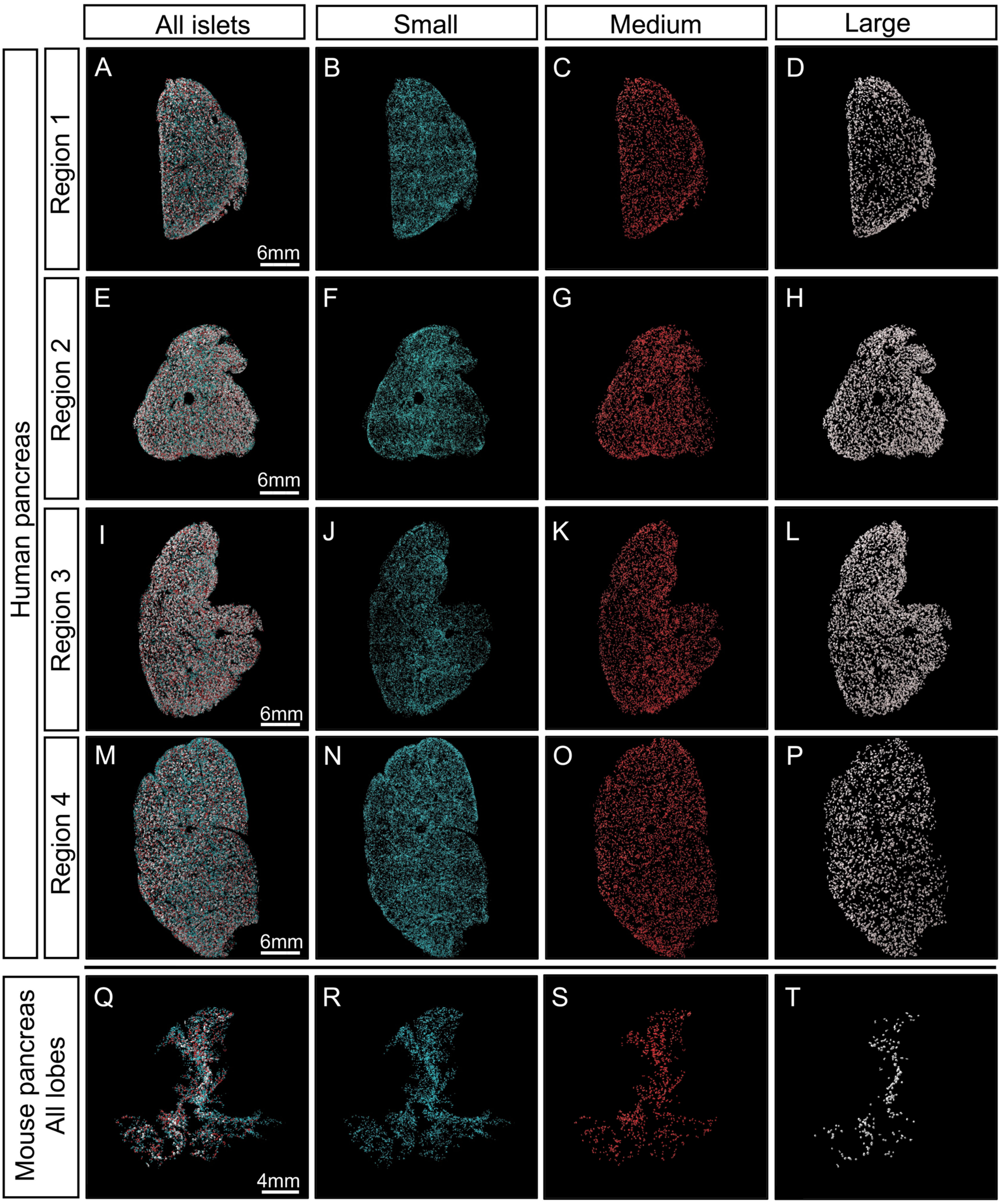


**Extended Data Fig. 6. Comparative distribution of islets of different size categories in human and mouse pancreas.** (**A-P**) Human pancreatic discs (H2456) from region 1-4 in which the insulin signal has been segmented and pseudo colored according to size as defined in **Fig. 3**. (**Q-R**) Representative mouse pancreas from a C57BL/6 mouse at 10 weeks segmented and pseudo colored in the same way. Human islets show a more homogenous size distribution within the organ compared to mice in which large islets are primarily located along the central axis.

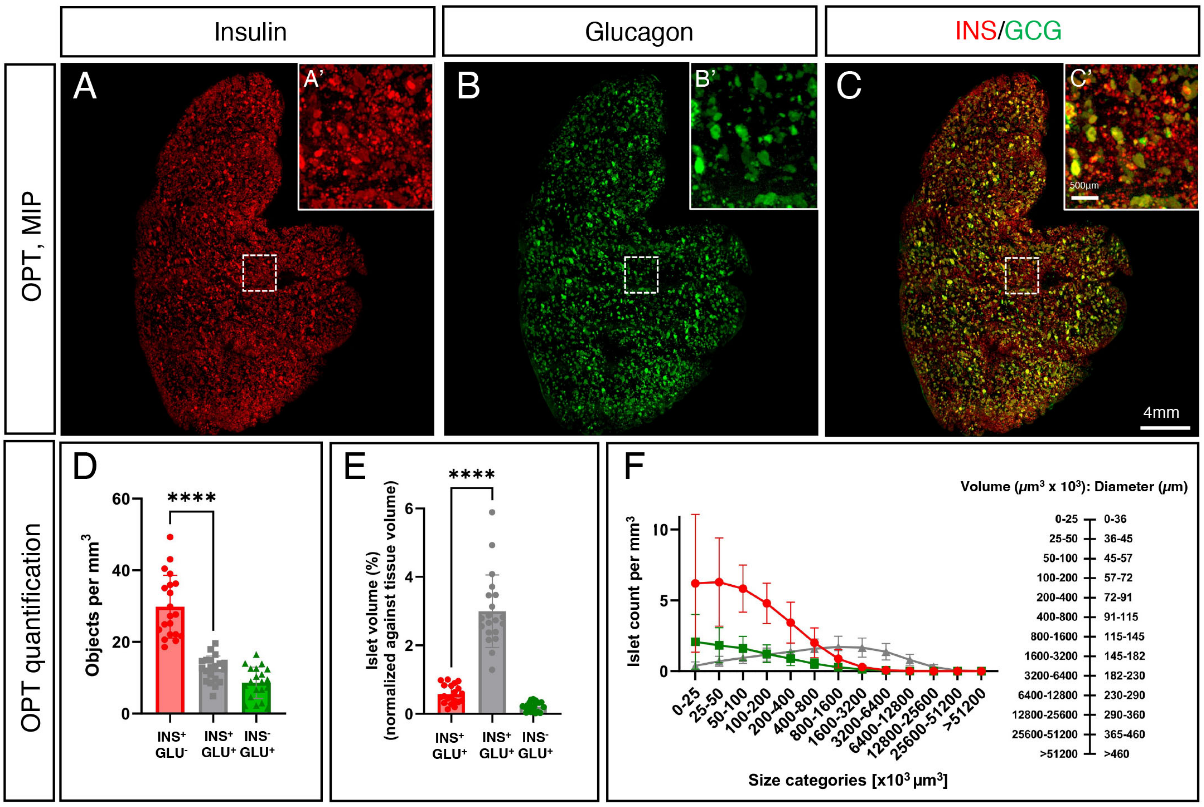


**Extended Data Fig. 7. OPT analyzes show that the majority of INS$^{+}$ islets are GCG$^{-}$.** (**A-C**) OPT maximum projection intensity (MIP) views of representative tissue discs from an ND donor (H2456, see **Extended Data Table 1**), showing insulin (**A**, red), glucagon (**B**, green) and both channels together (**C**). Note the substantial fraction of INS$^{+}$GCG$^{-}$ islets (see insets in panels **A-C**). (**D**) A graph showing the average densities of INS$^{+}$GCG$^{-}$, INS$^{-}$GCG$^{+}$ and INS$^{+}$GCG$^{+}$ islets from regions 1-4 in five donor pancreata (H2456, H2457, H2466, H2506 and H2522, **Extended Data Table 1,** see also **Extended Data Fig. 6**) encompassing a total of 824,154 islets. (**E**) A graph showing the volume constituted by each of the three categories displayed in **D** normalized to the total tissue volume. (**F**) Densities of all three islet subtypes (i.e., INS$^{+}$GCG$^{-}$, INS$^{-}$GCG$^{+}$ and INS$^{+}$GCG$^{+}$) in **E** represented as a function of islet size. Note that the majority of INS$^{+}$GCG$^{-}$ islets belong to the smaller size categories, with a volume typically < 200-400 x $10^3$ µm$^3$ (=72-91 µm in diameter). Error bars show mean ± SD. Scale bar in **C** is 4 mm (for main images in **A-C**) and scale bar in inset C´ is 500 µm (for A´-C´). **** donates a statistical significance of 0.0001.

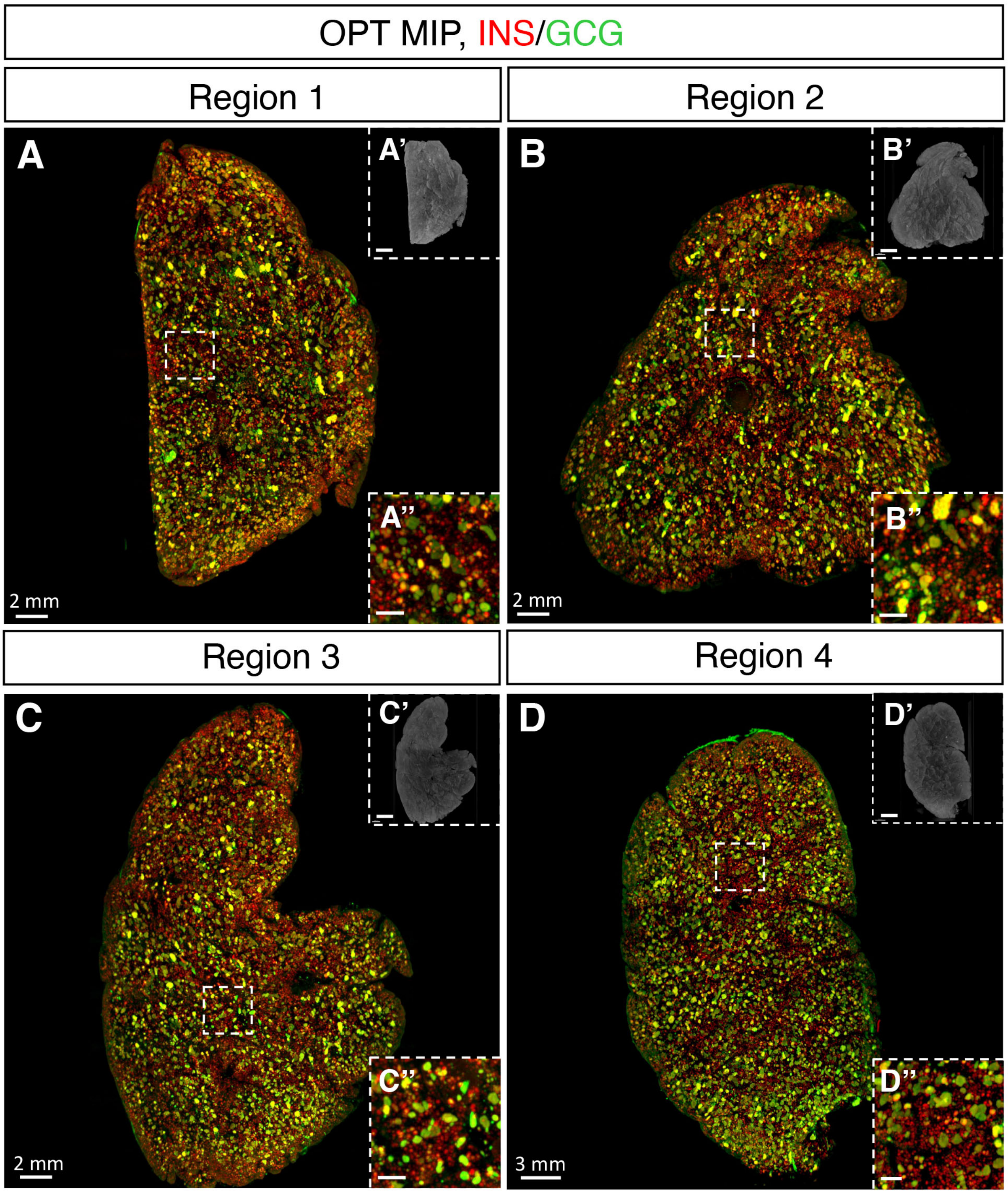


**Fig. S8. OPT images of INS and GCG in human pancreatic discs from regions 1-4.** (**A-D**) Representative OPT maximum intensity projection (MIP) images from scans from regions 1-4, showing insulin (INS, red) and glucagon (GCG, green). Top panel insets show AF disc anatomy of the areas within the dotted boxes, whereas bottom panel insets highlight smaller islets that merely express insulin. Scale bar in **A-D** is 500 μm in the inset (**A´´-D´´**).

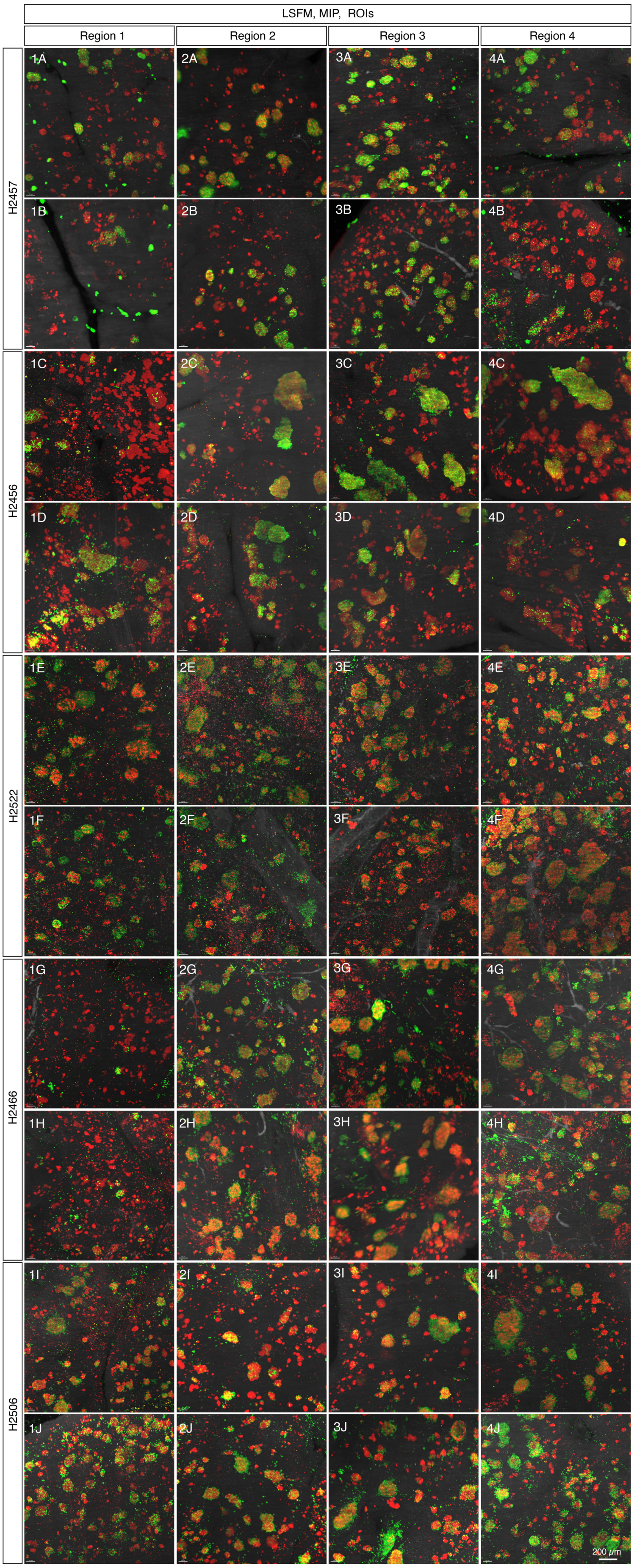


**Extended Data Fig. 9. LSFM analyzes of insulin (INS) and glucagon (GCG).** (**A-P**) LSFM analyzes of ROIs (sampled from H2456, H2457, H2466, H2506 and H2522, see **Extended Data table 1**) used for statistical assessment of islet heterogeneity (see **Fig. 4**) showing insulin (INS, red) and glucagon (GCG, green). Scan depth was 1 mm. Scale bar in **P** is 200 $\mu$m.

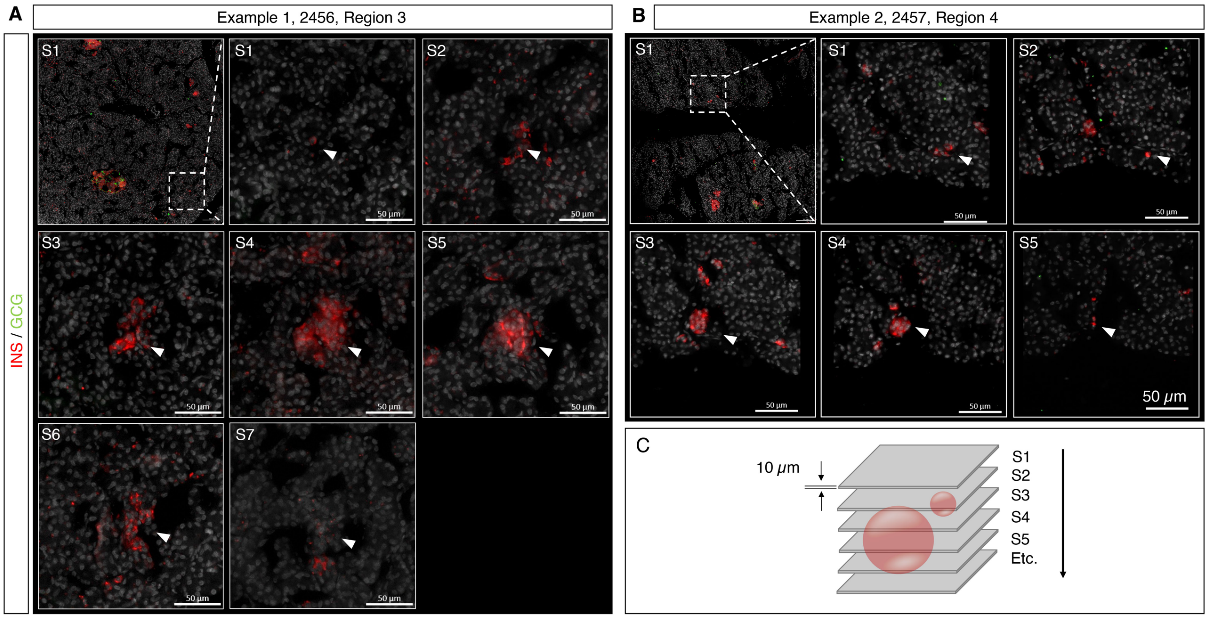


**Extended Data Fig. 10. Examples of Axioscan slide scanner analyzes demonstrating the absence of GCG$^{+}$ cells in INS$^{+}$ islets in 2D cryosections.** By assessing islets as a Z-stack section (S) by section (10 μm) throughout the islet volume (white arrow head), 3D optical data of INS$^{+}$GCG$^{-}$ islets were confirmed. Note the presence of GCG$^{+}$ cells (green) in the zoomed-out view in **A** (S1, arrow) and **B** (S1, arrow). Scale bar is 50 μm.

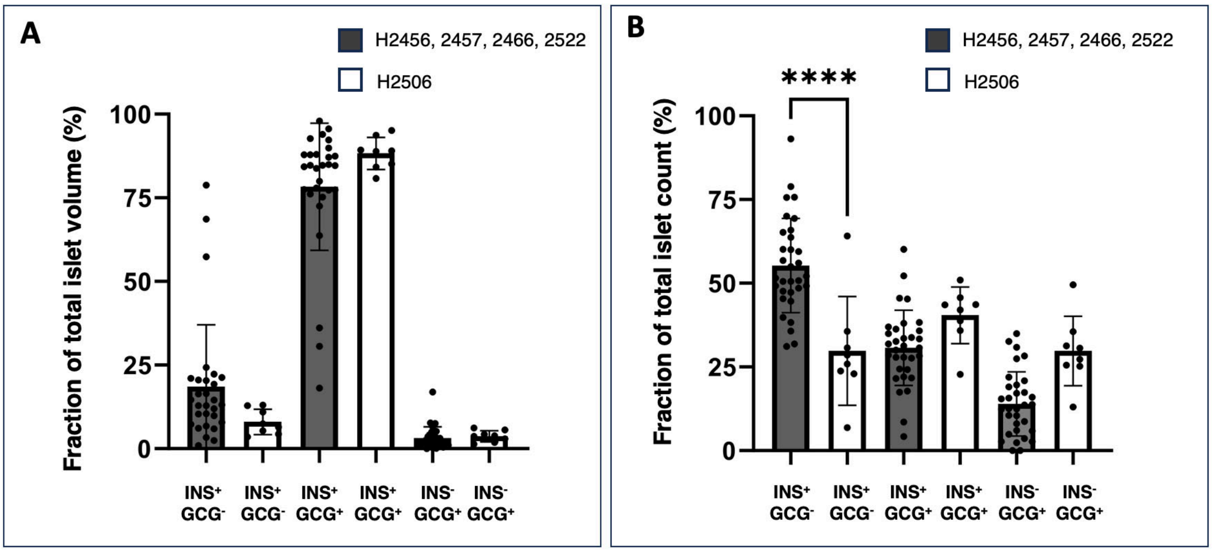


**Extended Data Fig. 11. Comparison of LSFM derived fractional statistics between H2506 and (H2456, 2457, 2466, 2522).** (**A**) Fraction of total islet volume and (**B**) fraction of total islet volume for each category. **** donates a statistical significance of 0.0001.

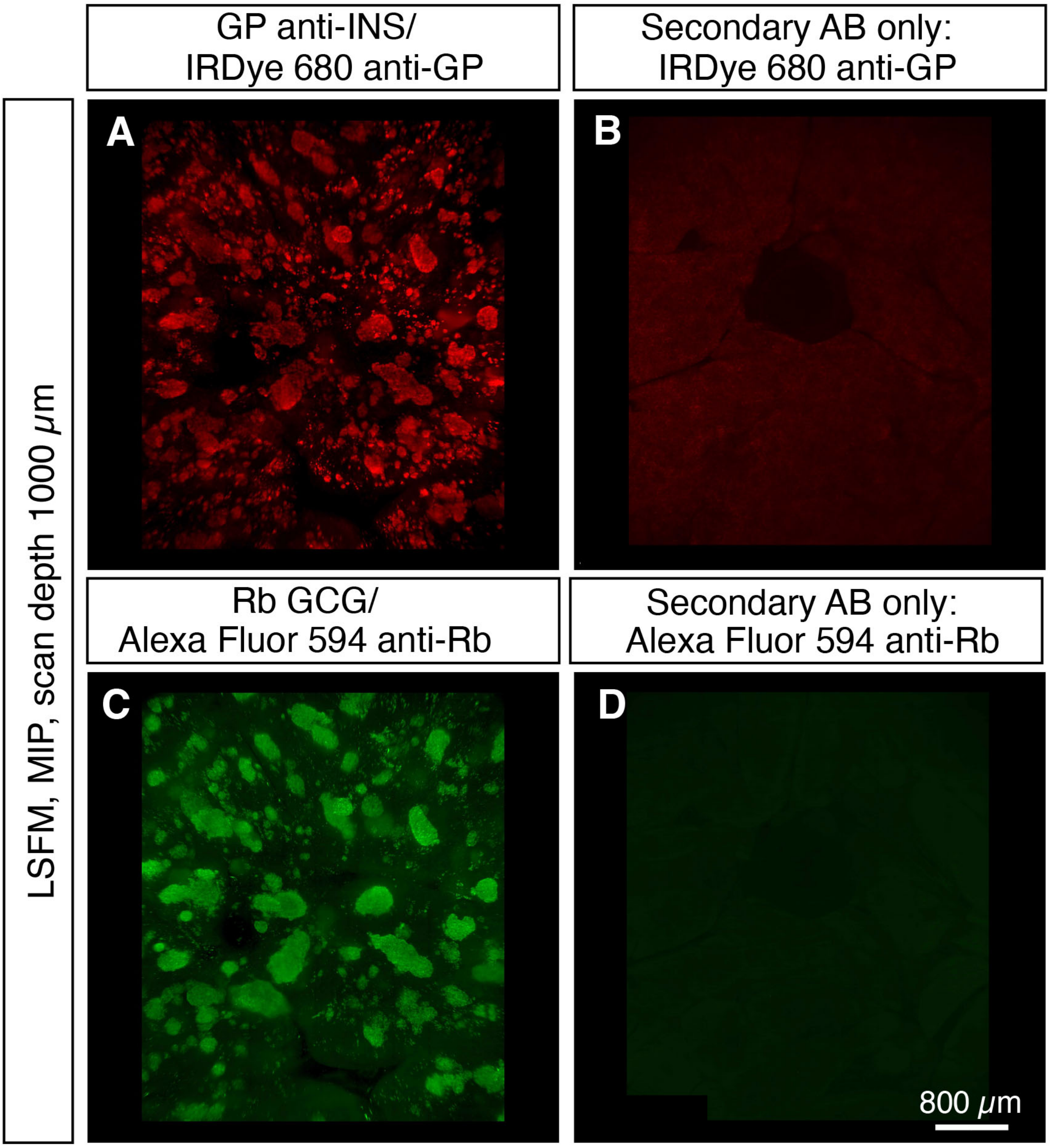


**Extended Data Fig. 12. Antibody controls.** LSFM datasets showing primary and secondary antibodies for staining (**A**) insulin (INS) and (**C**) glucagon (GCG), as well as and secondary antibodies only controls in panels **B** (anti-guinea-pig IgG, 680) and **D** (anti-rabbit IgG, 594). In all cases, antibody incubation times and scan settings were identical.

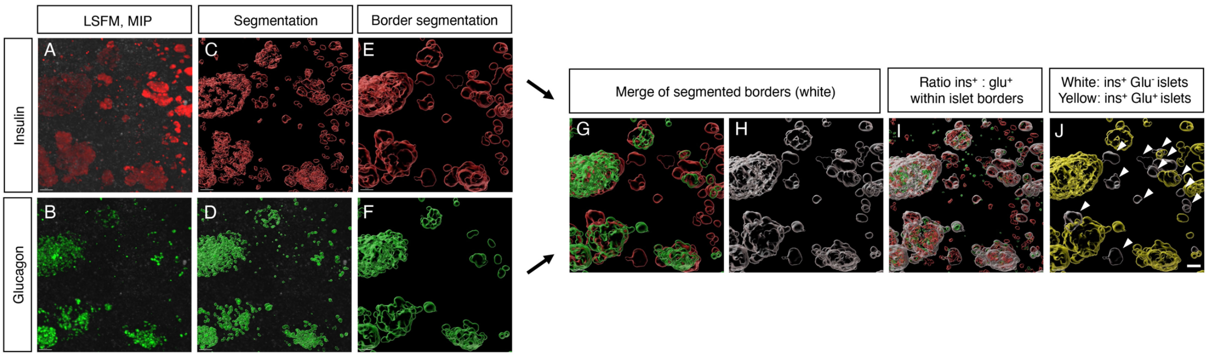


**Extended Data Fig. 13. Flow chart exemplifying the pipeline for LSFM-based assessment of INS:GCG ratios in human pancreatic islets.** (**A**, **B**) LSFM maximum intensity projection (MIP) view of INS (**A**, red) and GCG (**B**, green). (**C, D**) Segmentation of the INS$^+$ (**C**) and GCG$^+$ (**D**) channels, respectively. (**E, F**) Segmentation of the outer border of the respective channel. (**G, H**) Segmented border showed together (**G**) and fused (**H**, white). INS$^+$ and GCG$^+$ (red and green, respectively) within fused outer borders (white) for INS:GCG ratios. (**J)** Pseudo-coloring of segmented islet volumes >25 $\mu$m in Ø for which islets (INS$^+$ + GCG$^+$ outer border volume) containing <1% GCG$^+$ signals are pseudo-colored white and all other islets are coloured yellow. Note that the threshold was set to 1%, primarily to exclude any inclusion of general signal/labeling noise (see methods). Scale bar in (**J**) is 50 $\mu$m.

| Donor | Sex | Age | BMI | HbA1c | Figures | Comment | $INS^{+}GLU^{-}$ Islets (%) |
|---|---|---|---|---|---|---|---|
| H2456 | M | 26 | 31.7 | 29 | 3A-C, 4A-E, S6, S7, S8A, S9, S10 | Discs from reg 1-4 | 56,47 |
| H2457 | M | 28 | 23.7 | 35 | 1A-D, S1A-B, S2, S3, S7, S8B | Entire pancreas | 62,76 |
| H2466 | F | 38 | 37.6 | 38 | S7 | Discs from reg 1-4 | 51,69 |
| H2506 | M | 20 | 23.8 | 34 | S7 | Discs from reg 1-4 | 29.80 |
| H2522 | F | 45 | 25.1 | NA | S7 | Discs from reg 1-4 | 50.19 |

**Extended Data Table 1. Donor data and clinical parameters.**

| Specimen 1 | Manual annotation | Segmentation | Accuray (%) | Average accuracy (%) |
|---|---|---|---|---|
| ROI 1 | 962 | 1143 | 119 | 102.3 (SD: +/- 8.09) |
| ROI 2 | 791 | 778 | 98 | |
| ROI 3 | 727 | 724 | 100 | |
| **Specimen 2** | | | | |
| ROI 1 | 655 | 648 | 99 | |
| ROI 2 | 939 | 931 | 99 | |
| ROI 3 | 627 | 576 | 92 | |
| **Specimen 3** | | | | |
| ROI 1 | 671 | 706 | 105 | |
| ROI 2 | 760 | 746 | 98 | |
| ROI 3 | 937 | 1038 | 111 | |

**Extended Data Table 2. Evaluation of accuracy between manual annotation and computational segmentation.**
