## Supplementary material for "Illuminating the complete ß-cell mass of the human pancreas - signifying a new view on the islets of Langerhans": Movie legends Movie S1-S5

**Movie S1**. Maximum projection intensity (MIP) of a pancreatic disc (H2457) showing insulin labeled islets (red) and anatomical outline based on tissue autofluoresecence (grey).

**Movie S2**. Maximum projection intensity (MIP) of combined NIR-OPT datasets displaying the complete β-cell mass distribution of a representative human pancreas. The displayed pancreas (H2457) contains 1.17 $cm^3$ $INS^+$ cells comprising 2.21 x $10^6$ $INS^+$ islets. Note, due to size limitations the movie is significantly down sized.

**Movie S3**. Movie showing a disc from a representative human donor pancreas in which the insulin signal has been segmented and pseudo colored according to size. Each size category (blue, small; red, medium; white, large) corresponds to 1/3 of the total β-cell volume of the pancreas. Note, due to size limitations the movie is significantly down sized.

**Movie S4**. Movie showing a representative pancreas from a C57Bl/6 mouse at 10 weeks in which the insulin signal has been segmented and pseudo colored according to size. Each size category (blue, small; red, medium; white, large) corresponds to 1/3 of the total β-cell volume in 5 animals. In mice, large islets are predominantly distributed along the central axis following the main pancreatic duct. Note, due to size limitations the movie is significantly down sized.

**Movie S5**. Maximum intensity projection (MIP) from a LSFM scan of a representative ROI (see methods) from a ND donor pancreas stained for insulin (INS, red) and glucagon (GCG, green) showing the presence of $INS^+GCG^-$ islets.
